## Supplementary Figures for "Low dose rate γ-irradiation protects *Drosophila melanogaster* chromosomes from double strand breaks and telomere fusions by modulating the expression of the esi-RNA biogenesis factor *Loquacious*"

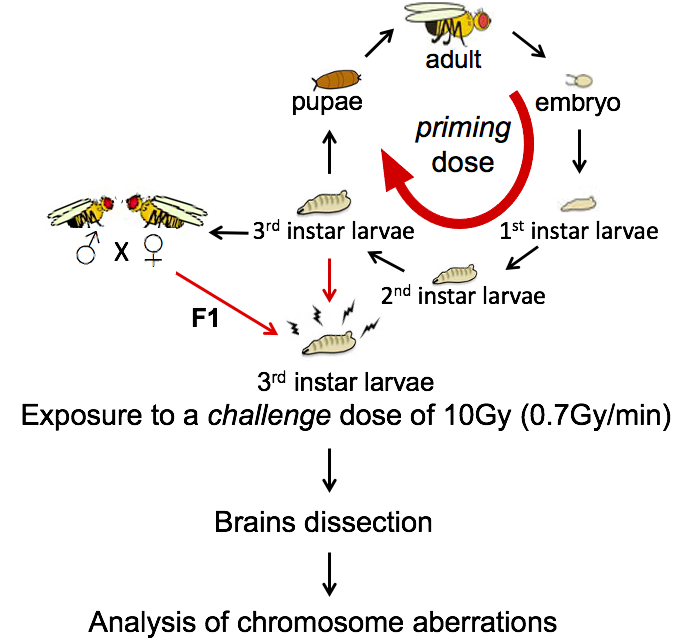

**Supplementary Figure 1.** Schematic representation of the experimental set up for the analysis of radioresistance. The *priming* dose of 0.4Gy (LDR) was delivered with a dose-rate of 2.5mGy/h during the embryos-larvae transition. The third instar larvae were collected and exposed to the *challenge* dose of 10Gy delivered with a dose-rate of 0.7Gy/min. Brains were then dissected and analyzed for the Chromosome aberrations frequency. Males and females pretreated (0.4Gy LDR) during their development were crossed to obtain F_1_. The third instar F_1_ larvae were exposed to the *challenge* dose of 10Gy to analyze the Chromosome aberrations frequency.

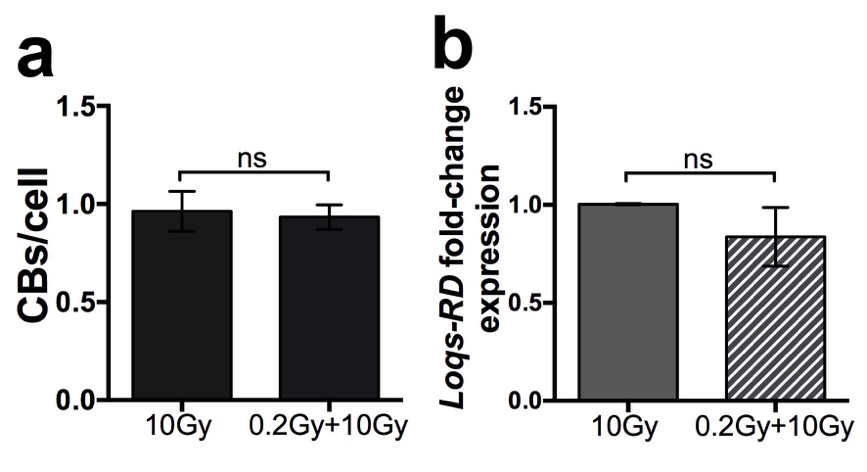

**Supplementary Figure 2. A protracted 0.2Gy (1.25mGy/h) g dose does neither trigger RAR nor reduce *Loqs*-RD transcripts (a)** Frequencies of CBs in both not-treated and 0.2Gy (1.25mGy/h) irradiated *Oregon*-R larval brains after the exposure to the *challenging* dose of 10Gy**.** CBs: chromosome breaks; ns: no significant. At least three independent experiments were conducted, with 200 cells analyzed. The statistical analysis was assessed using the Student t-test. **(b)** RT-qPCR results of *Loqs-RD* performed in samples chronically exposed to a low dose of 0.2Gy and then treated with 10Gy compared to samples exposed to 10Gy alone. The statistical analysis was conducted using the Student *t-test*; ns: not significant.

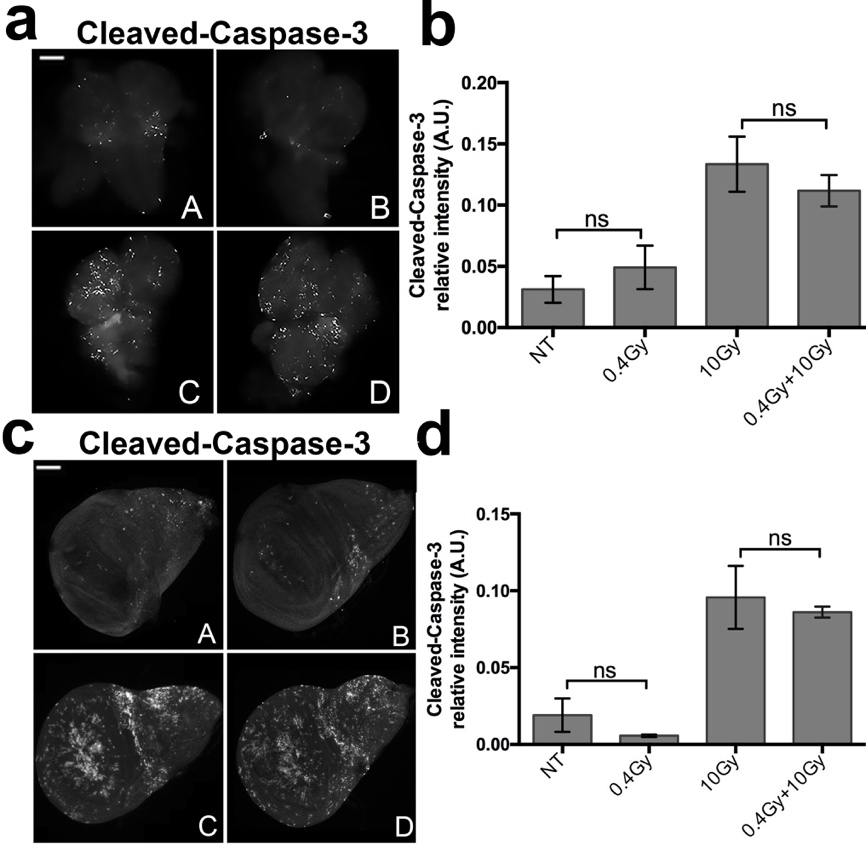

**Supplementary Figure 3. Apoptosis analysis in irradiated larval brains and wing imaginal discs.(a)** Immunofluorescence with anti-cleaved-caspase-3 on larval brains; (A, C) Untreated and (B, D) 0.4Gy LDR disc after exposure to 10Gy. **(b)** Quantification of Cleaved-Caspase-3 signals; ns: no statistically significant values; NT: non-treated. **(c)** Immunofluorescence with anti-cleaved-caspase-3 on wing imaginal discs; (A, C) Untreated and (B, D) 0.4Gy LDR disc after exposure to 10Gy. **(d)** Quantification of Cleaved-Caspase-3 signals; ns: no statistically significant values; NT: non-treated.

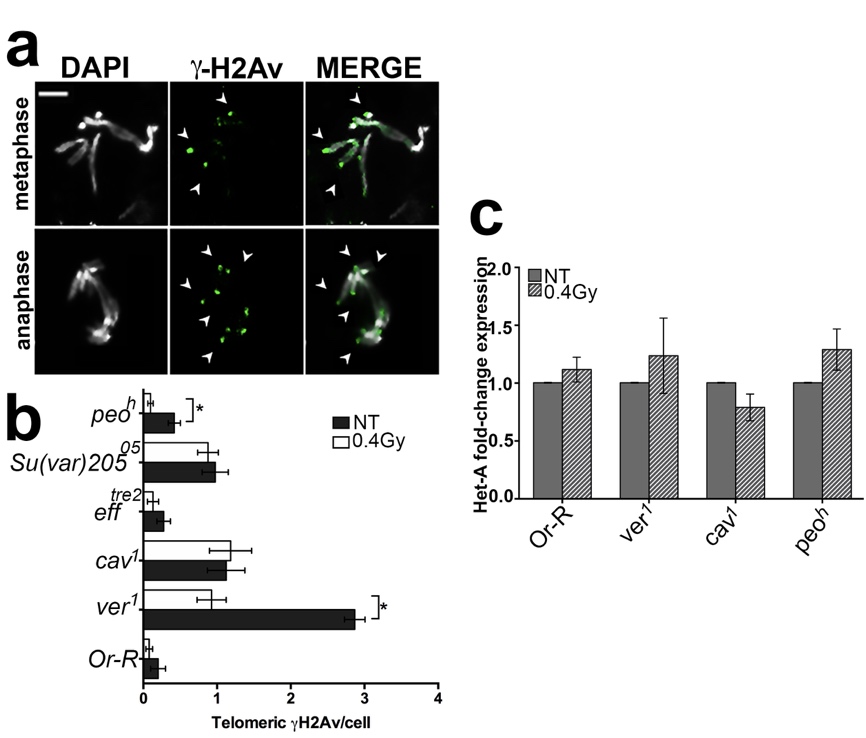

**Supplementary Figure 4. Telomeric γH2Av foci analysis and Het-A expression in 0.4Gy LDR telomere capping mutants. (a)**. Examples of γH2Av localization (green) on chromosome ends (arrows) from a mutant metaphase and anaphase. **(b)** Frequency of Telomeric γH2Av foci in 0.4Gy LDR telomere capping mutants. Student t-test (**p*<0.05); NT: non-treated. **(c)** Het-A expression in 0.4Gy LDR and untreated *Or*-R, *ver^1^*, *cav^1^*, and *peo^h^* third instar larvae. The qPCR shows no statistically significative differences between the 0.4Gy LDR-treated samples and the corresponding untreated controls. Student t-test (*p*<0.05); NT: non-treated.

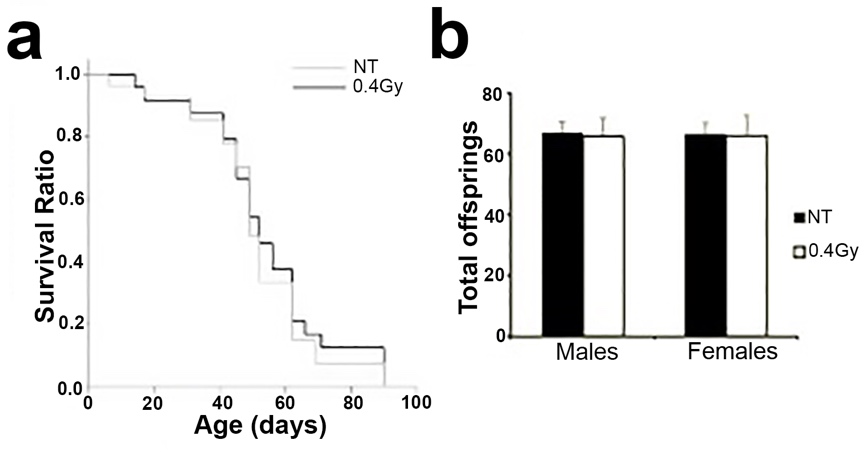

**Supplementary figure 5. 0.4Gy-LDR ionizing radiation doses do not affect *Oregon*-R life span and fertility. a)** Life span Kaplan-Meier analysis of *Oregon*-R males treated (0.4Gy LDR, black line) and non-treated (NT, gray line). The survival ratio is indistinguishable in the two samples. Long-rank: 0.249; p=0.618. **(b)** The low dose exposure does not influence the male and female fertility. In the graph are represented the average number of individuals born from three different crosses. NT: non-treated.
