## Supplementary Tables for "Low dose rate γ-irradiation protects *Drosophila melanogaster* chromosomes from double strand breaks and telomere fusions by modulating the expression of the esi-RNA biogenesis factor *Loquacious*"

**Supplementary Table 1**. **Frequency of chromosome breaks observed in F_1_ and F_2_ generations from 0.4Gy LDR-treated parents.**

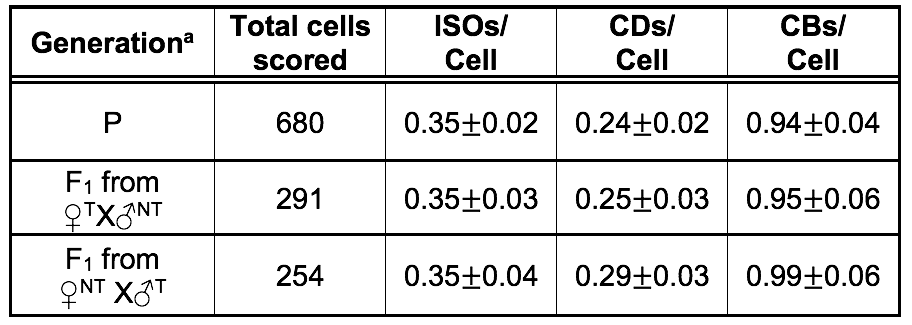

Quantification of CBs in F_1_ and F_2_ generations obtained from *Oregon-R* parents exposed, during their embryo-third instar larvae transition, to a 0.4Gy LDR γ ray treatment. Note that, following the 10Gy acute exposure, CBs frequency is reduced in F_1_ but not in F_2_.. The statistical analysis was performed using the t-Student Test. a= Third instar larvae exposed to a 10Gy *challenge* dose have been analyzed for each generation; P: parental generation; F_1_: first filial generation; F_2_: second filial generation; T: treated; NT: non-treated; ISOs: isochromosome deletions; CD: chromatid deletions; CBs: chromosome breaks.

**Supplementary Table 2**. **Frequency of chromosome breaks observed in *nbs^1^* single mutants and *loqs^KO^;nbs^1^* double mutants.**

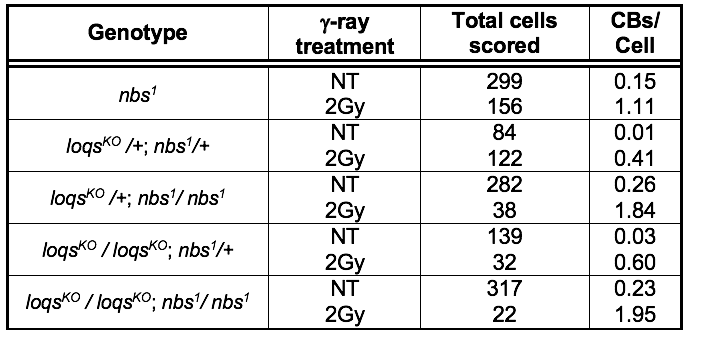

Frequency of IR-induced CBs in *nbs*^1^ single mutant and *loqs*;*nbs^1^* double mutants. Note that loss of Nbs abolishes the radioresistance observed in *loqs^KO^* mutants. See text for details. The statistical analysis performed was the *Student t-test*. CBs: chromosome breaks. NT: non-treated

**Supplementary Table 3**. **Differentially expressed genes between 0.4 Gy LDDR + 10 Gy-irradiated *vs*. unirradiated larval male brains.**

A list of 106 differentially expressed genes (94 up-regulated and 12 down-regulated) from 0.4 Gy LDDR + 10 Gy *vs*. unirradiated (D *vs*. A) comparison. considering a 5% false discovery rate. In the last column are represented the expression levels of each transcript calculated as Log2(D/A).

| **FlyBase ID** | **Enrez ID** | **Description** | **log2(D/A)** |
| --- | --- | --- | --- |
| agt | 40816 | O-6-alkylguanine-DNA alkyltransferase(agt) | **0.62** |
| Amnionless | 33199 | nogaster Amnionless ortholog(Amnionless) | **0.64** |
| AOX1 | 41894 | Aldehyde oxidase 1(AOX1) | **1.33** |
| CG10005 | 41451 | CG10005 gene product from transcript CG10005-RA(CG10005) | **0.76** |
| CG10638 | 39424 | CG10638 gene product from transcript CG10638-RD(CG10638) | **0.66** |
| CG11131 | 40511 | CG11131 gene product from transcript CG11131-RB(CG11131) | **0.37** |
| CG11897 | 43450 | CG11897 gene product from transcript CG11897-RB(CG11897) | **1.01** |
| CG12171 | 40690 | CG12171 gene product from transcript CG12171-RA(CG12171) | **0.53** |
| CG12224 | 41453 | CG12224 gene product from transcript CG12224-RC(CG12224) | **0.63** |
| CG12264 | 34613 | CG12264 gene product from transcript CG12264-RA(CG12264) | **0.72** |
| CG1299 | 38496 | CG1299 gene product from transcript CG1299-RA(CG1299) | **0.65** |
| CG13067 | 39785 | CG13067 gene product from transcript CG13067-RA(CG13067) | **0.55** |
| CG13705 | 38587 | CG13705 gene product from transcript CG13705-RA(CG13705) | **0.40** |
| CG14419 | 31276 | CG14419 gene product from transcript CG14419-RA(CG14419) | **0.38** |
| CG15784 | 31461 | CG15784 gene product from transcript CG15784-RB(CG15784) | **1.12** |
| CG1582 | 32021 | CG1582 gene product from transcript CG1582-RB(CG1582) | **0.51** |
| CG17104 | 34465 | CG17104 gene product from transcript CG17104-RB(CG17104) | **0.67** |
| CG1809 | 35981 | CG1809 gene product from transcript CG1809-RA(CG1809) | **0.00** |
| CG18547 | 41452 | CG18547 gene product from transcript CG18547-RA(CG18547) | **1.48** |
| CG3008 | 33693 | CG3008 gene product from transcript CG3008-RA(CG3008) | **0.85** |
| CG31869 | 34490 | CG31869 gene product from transcript CG31869-RC(CG31869) | **0.59** |
| CG32071 | 39246 | CG32071 gene product from transcript CG32071-RA(CG32071) | **0.24** |
| CG32603 | 318109 | CG32603 gene product from transcript CG32603-RA(CG32603) | **0.18** |
| CG33158 | 39834 | CG33158 gene product from transcript CG33158-RB(CG33158) | **0.51** |
| CG34417 | 31591 | CG34417 gene product from transcript CG34417-RI(CG34417) | **0.39** |
| CG3448 | 39094 | CG3448 gene product from transcript CG3448-RB(CG3448) | **0.78** |
| CG4115 | 41499 | CG4115 gene product from transcript CG4115-RA(CG4115) | **0.80** |
| CG42326 | 35838 | CG42326 gene product from transcript CG42326-RE(CG42326) | **0.63** |
| CG43427 | 40583 | CG43427 gene product from transcript CG43427-RP(CG43427) | **0.33** |
| CG43693 | 39231 | CG43693 gene product from transcript CG43693-RD(CG43693) | **-0.41** |
| CG5059 | 40278 | CG5059 gene product from transcript CG5059-RD(CG5059) | **-0.32** |
| CG5205 | 41891 | CG5205 gene product from transcript CG5205-RA(CG5205) | **0.55** |
| CG5955 | 40268 | CG5955 gene product from transcript CG5955-RA(CG5955) | **0.74** |
| CG6512 | 39922 | CG6512 gene product from transcript CG6512-RB(CG6512) | **0.45** |
| CG6901 | 41977 | CG6901 gene product from transcript CG6901-RA(CG6901) | **0.61** |
| CG7627 | 34148 | CG7627 gene product from transcript CG7627-RB(CG7627) | **0.53** |
| CG8064 | 42196 | CG8064 gene product from transcript CG8064-RA(CG8064) | **0.30** |
| CG9090 | 37297 | CG9090 gene product from transcript CG9090-RA(CG9090) | **0.58** |
| CG9297 | 41688 | CG9297 gene product from transcript CG9297-RC(CG9297) | **0.62** |
| CG9411 | 32392 | CG9411 gene product from transcript CG9411-RA(CG9411) | **0.61** |
| CG9759 | 41661 | CG9759 gene product from transcript CG9759-RB(CG9759) | **0.40** |
| CHKov2 | 43068 | CG10675 gene product from transcript CG10675-RA(CHKov2) | **0.68** |
| Cht6 | 31935 | CG43374 gene product from transcript CG43374-RK(Cht6) | **0.55** |
| Corp | 31764 | Companion of reaper(Corp) | **0.66** |
| Cpr100A | 43657 | Cuticular protein 100A(Cpr100A) | **0.43** |
| CR43144 | 19834780 | ncRNA(CR43144) | **0.74** |
| CR44272 | 19835298 | ncRNA(CR44272) | **-0.61** |
| CR44430 | 19835618 | ncRNA(CR44430) | **0.70** |
| CR44751 | 19834965 | ncRNA(CR44751) | **0.46** |
| CR46003 | 26067324 | ncRNA(CR46003) | **-0.32** |
| cv-2 | 45280 | crossveinless 2(cv-2) | **0.80** |
| dally | 39013 | division abnormally delayed(dally) | **0.32** |
| Dif | 35045 | Dorsal-related immunity factor(Dif) | **0.46** |
| dp | 36461 | DP transcription factor(Dp) | **0.41** |
| dyl | 38531 | dusky-like(dyl) | **0.92** |
| E(spl)m5-HLH | 43158 | Enhancer of split m5. helix-loop-helix(E(spl)m5-HLH) | **-0.63** |
| E(spl)m6-BFM | 43159 | Enhancer of split m6. Bearded family member(E(spl)m6-BFM) | **-0.59** |
| E(spl)mgamma-HLH | 43151 | Enhancer of split mgamma. helix-loop-helix(E(spl)mgamma-HLH) | **-0.60** |
| Ect4 | 38895 | Ectoderm-expressed 4(Ect4) | **0.36** |
| egr | 36054 | eiger(egr) | **0.35** |
| form3 | 3346238 | formin 3(form3) | **0.35** |
| Gadd45 | 35646 | CG11086 gene product from transcript CG11086-RA(Gadd45) | **0.84** |
| Gclc | 53581 | Glutamate-cysteine ligase catalytic subunit(Gclc) | **0.50** |
| Gcn5 | 39431 | Gcn5 ortholog(Gcn5) | **0.32** |
| GstD3 | 48336 | Glutathione S transferase D3(GstD3) | **0.56** |
| GstE5 | 37110 | Glutathione S transferase E5(GstE5) | **0.60** |
| GstE6 | 37111 | Glutathione S transferase E6(GstE6) | **0.95** |
| hid | 40009 | head involution defective(hid) | **0.89** |
| Hmgcr | 42803 | HMG Coenzyme A reductase(Hmgcr) | **0.47** |
| HmgZ | 37480 | HMG protein Z(HmgZ) | **-0.44** |
| Hsc70-5 | 36583 | Heat shock protein cognate 5(Hsc70-5) | **0.29** |
| Hsp60 | 32045 | Heat shock protein 60(Hsp60) | **0.36** |
| Ilp8 | 39909 | Insulin-like peptide 8(Ilp8) | **1.28** |
| Irbp | 117419 | Inverted repeat-binding protein(Irbp) | **0.72** |
| JhI-26 | 36819 | Juvenile hormone-inducible protein 26(JhI-26) | **0.70** |
| Ku80 | 34930 | CG18801 gene product from transcript CG18801-RA(Ku80) | **0.67** |
| l(3)72Ab | 39737 | lethal (3) 72Ab(l(3)72Ab) | **0.31** |
| magu | 36048 | CG2264 gene product from transcript CG2264-RA(magu) | **-0.47** |
| Mdr49 | 36428 | Multi drug resistance 49(Mdr49) | **0.70** |
| mfas | 41455 | midline fasciclin(mfas) | **0.27** |
| Mocs1 | 39238 | Molybdenum cofactor synthesis 1 ortholog(Mocs1) | **0.59** |
| mre11 | 34565 | meiotic recombination 11(mre11) | **0.55** |
| MRP | 34686 | Multidrug-Resistance like Protein 1(MRP) | **0.75** |
| mus205 | 47186 | mutagen-sensitive 205(mus205) | **0.54** |
| nerfin-1 | 44786 | nervous fingers 1(nerfin-1) | **-0.21** |
| Nplp4 | 50190 | Neuropeptide-like precursor 4(Nplp4) | **1.09** |
| PEK | 40653 | pancreatic eIF-2alpha kinase(PEK) | **0.35** |
| pic | 41611 | piccolo(pic) | **0.27** |
| POSH | 36990 | Plenty of SH3s(POSH) | **0.30** |
| Pvf2 | 33994 | PDGF- and VEGF-related factor 2(Pvf2) | **0.49** |
| RnrL | 34392 | Ribonucleoside diphosphate reductase large subunit(RnrL) | **0.34** |
| roX1 | 3772376 | RNA on the X 1(roX1) | **-0.31** |
| rpr | 40015 | reaper(rpr) | **0.73** |
| scaf | 35505 | scarface(scaf) | **0.61** |
| scb | 36692 | scab(scb) | **0.65** |
| sda | 44359 | slamdance(sda) | **0.36** |
| snoRNA:Psi18S-525c | 5740385 | ncRNA(snoRNA:Psi18S-525c) | **-0.58** |
| spn-E | 41919 | spindle E(spn-E) | **0.85** |
| Sulf1 | 53437 | Sulfated(Sulf1) | **0.56** |
| Traf4 | 33638 | TNF-receptor-associated factor 4(Traf4) | **0.43** |
| Ugt86Da | 53510 | CG18578 gene product from transcript CG18578-RA(Ugt86Da) | **0.79** |
| Ugt86Di | 53502 | CG6658 gene product from transcript CG6658-RB(Ugt86Di) | **0.94** |
| Vinc | 31201 | Vinculin(Vinc) | **0.38** |
| wnd | 40143 | wallenda(wnd) | **0.27** |
| Xrp1 | 42267 | CG17836 gene product from transcript CG17836-RB(Xrp1) | **0.53** |
| zip | 38001 | zipper(zip) | **0.31** |

**Supplementary Table 4.** **Differentially expressed genes between 0.4 Gy LDDR + 10 Gy *vs*. 10 Gy-irradiated larval male brains.**

A list of 107 differentially expressed genes (42 up-regulated and 65 down-regulated) from 0.4 Gy LDDR + 10 Gy *vs*. 10 Gy (D *vs*. B) comparison. considering a 5% false discovery rate. In the last column are represented the expression levels of each transcript calculated as Log_2_ (D/B).

| **FlyBase ID** | **Entrez ID** | **Description** | **Log2(D/B)** |
| --- | --- | --- | --- |
| AATS-ILE | 45785 | Isoleucyl-tRNA synthetase [Source:FlyBase | **0.32** |
| ALDH | 34256 | Aldehyde dehydrogenase [Source:FlyBase | **-0.43** |
| AMNIONLESS | 33199 | Amnionless ortholog [Source:FlyBase | **0.51** |
| AOX1 | 41894 | Aldehyde oxidase 1 [Source:FlyBase | **0.48** |
| AURB | 34504 | aurora B [Source:FlyBase | **-0.51** |
| BEAT-IIIC | 35037 | beat-IIIc [Source:FlyBase | **-0.34** |
| BWA | 250736 | brain washing [Source:FlyBase | **-0.39** |
| CAND1 | 34403 | Cullin-associated and neddylation-dissociated 1 [Source:FlyBase | **0.28** |
| CG10005 | 41451 | RE59626p [Source:UniProtKB/TrEMBL | **0.54** |
| CG11151 | 32335 | GEO07753p1 [Source:UniProtKB/TrEMBL | **-0.49** |
| CG12194 | 33685 | Uncharacterized protein. isoform B [Source:UniProtKB/TrEMBL | **-0.28** |
| CG12325 | 36144 | LD10780p [Source:UniProtKB/TrEMBL | **0.40** |
| CG12499 | 42637 | NA | **0.37** |
| CG13185 | 36268 | NA | **0.39** |
| CG13784 | 34003 | Uncharacterized protein. isoform E [Source:UniProtKB/TrEMBL | **-0.29** |
| CG17680 | 37071 | Essential MCU regulator. mitochondrial [Source:UniProtKB/Swiss-Prot | **-0.39** |
| CG1896 | 43738 | CG1896 [Source:UniProtKB/TrEMBL | **-0.44** |
| CG30349 | 35885 | FI03455p [Source:UniProtKB/TrEMBL | **0.32** |
| CG31683 | 261623 | BcDNA.GH02384 [Source:UniProtKB/TrEMBL | **-0.55** |
| CG31869 | 34490 | Uncharacterized protein. isoform A [Source:UniProtKB/TrEMBL | **0.40** |
| CG32026 | 317829 | GH12815p [Source:UniProtKB/TrEMBL | **0.43** |
| CG32318 | 2768976 | RT07324p [Source:UniProtKB/TrEMBL | **-0.52** |
| CG32344 | 326208 | LD28101p [Source:UniProtKB/TrEMBL | **0.28** |
| CG33158 | 39834 | NA | **0.42** |
| CG34159 | 34353 | GEO11246p1 [Source:UniProtKB/TrEMBL | **-0.43** |
| CG34232 | 5740195 | LP22624p [Source:UniProtKB/TrEMBL | **-0.62** |
| CG3835 | 31184 | CG3835. isoform A [Source:UniProtKB/TrEMBL | **-0.49** |
| CG40228 | 5740664 | Transcription elongation factor 1 homolog [Source:UniProtKB/Swiss-Prot | **-0.37** |
| CG41128 | 3355108 | MICOS complex subunit MIC10 [Source:UniProtKB/TrEMBL | **-0.55** |
| CG42231 | 7354432 | NA | **-0.58** |
| CG42362 | 7354407 | IP15503p1 [Source:UniProtKB/TrEMBL | **0.54** |
| CG43349 | 12797873 | Uncharacterized protein. isoform A [Source:UniProtKB/TrEMBL | **-0.16** |
| CG4554 | 37570 | CG4554 [Source:UniProtKB/TrEMBL | **0.40** |
| CG5205 | 41891 | CG5205 [Source:UniProtKB/TrEMBL | **0.39** |
| CG6388 | 34656 | Probable tRNA (guanine(26)-N(2))-dimethyltransferase [Source:UniProtKB/Swiss-Prot | **0.38** |
| CG7607 | 39263 | CG7607 [Source:UniProtKB/TrEMBL | **-0.48** |
| CG7656 | 39691 | CG7656. isoform D [Source:UniProtKB/TrEMBL | **-0.35** |
| CG8064 | 42196 | CG8064 [Source:UniProtKB/TrEMBL | **0.36** |
| CG8939 | 32568 | Putative rRNA methyltransferase [Source:UniProtKB/TrEMBL | **0.47** |
| CLBN | 43018 | Caliban [Source:FlyBase | **0.47** |
| CR33987 | 3885608 | ncRNA(CR33987) | **-0.58** |
| CR43900 | 14462580 | ncRNA(CR43900) | **-0.65** |
| CR44272 | 19835298 | ncRNA(CR44272) | **-0.56** |
| CR45168 | 19835216 | ncRNA(CR45168) | **-0.75** |
| CR45820 | 26067148 | ncRNA(CR45820) | **0.58** |
| CR45908 | 26067236 | ncRNA(CR45908) | **-0.62** |
| CTPSYN | 39645 | CTP synthase [Source:FlyBase | **0.32** |
| CV-2 | 45280 | crossveinless 2 [Source:FlyBase | **0.40** |
| DEN1 | 36339 | Deneddylase 1 [Source:FlyBase | **-0.29** |
| DNAJ-60 | 37869 | DnaJ-like-60 [Source:FlyBase | **-0.57** |
| DRAT | 35687 | Death resistor Adh domain containing target [Source:FlyBase | **-0.37** |
| EIF4G | 43839 | eukaryotic translation initiation factor 4G [Source:FlyBase | **0.29** |
| GS1 | 33172 | Glutamine synthetase 1 [Source:FlyBase | **-0.35** |
| HEY | 35764 | Hairy/E(spl)-related with YRPW motif [Source:FlyBase | **-0.40** |
| HIS2B:CG33868 | 3772265 | His2B:CG33868 [Source:FlyBase | **0.23** |
| IFC | 33836 | infertile crescent [Source:FlyBase | **-0.27** |
| KRA | 40680 | krasavietz [Source:FlyBase | **0.43** |
| KUG | 40191 | kugelei [Source:FlyBase | **0.34** |
| L(2)35DF | 48782 | lethal (2) 35Df [Source:FlyBase | **0.30** |
| L(2)K09022 | 33960 | lethal (2) k09022 [Source:FlyBase | **0.40** |
| L(3)72AB | 39737 | lethal (3) 72Ab [Source:FlyBase | **0.28** |
| LK6 | 44672 | Lk6 [Source:FlyBase | **-0.25** |
| LOQS | 34751 | loquacious [Source:FlyBase | **-0.48** |
| MED7 | 41288 | Mediator complex subunit 7 [Source:FlyBase | **-0.50** |
| MPC1 | 42268 | Mitochondrial pyruvate carrier [Source:FlyBase | **-0.46** |
| MRP | 34686 | Multidrug-Resistance like Protein 1 [Source:FlyBase | **0.42** |
| MRPL33 | 50381 | mitochondrial ribosomal protein L33 [Source:FlyBase | **-0.50** |
| MRPS21 | 318249 | mitochondrial ribosomal protein S21 [Source:FlyBase | **-0.61** |
| MRPS28 | 37136 | mitochondrial ribosomal protein S28 [Source:FlyBase | **-0.49** |
| MYS45A | 35925 | Mystery 45A [Source:FlyBase | **0.39** |
| NPLP4 | 50190 | Neuropeptide-like precursor 4 [Source:FlyBase | **0.69** |
| PIC | 41611 | piccolo [Source:FlyBase | **0.25** |
| PINK1 | 31607 | PTEN-induced putative kinase 1 [Source:FlyBase | **-0.38** |
| PIS | 32506 | Phosphatidylinositol synthase [Source:FlyBase | **-0.40** |
| POSH | 36990 | Plenty of SH3s [Source:FlyBase | **0.34** |
| PTR | 35546 | Patched-related [Source:FlyBase | **-0.47** |
| RCA1 | 33959 | Regulator of cyclin A1 [Source:FlyBase | **-0.42** |
| RHEB | 117332 | Ras homolog enriched in brain ortholog (H. sapiens) [Source:FlyBase | **-0.38** |
| RME-8 | 35939 | Receptor mediated endocytosis 8 [Source:FlyBase | **0.26** |
| RNASEP:RNA | 3772418 | Ribonuclease P RNA [Source:FlyBase | **-0.56** |
| ROC2 | 36246 | Regulator of cullins 2 [Source:FlyBase | **-0.45** |
| RPI1 | 36617 | RNA polymerase I subunit [Source:FlyBase | **0.34** |
| RPL35 | 31483 | Ribosomal protein L35 [Source:FlyBase | **-0.46** |
| RPL36 | 31009 | Ribosomal protein L36 [Source:FlyBase | **-0.50** |
| RPL36A | 34098 | Ribosomal protein L36A [Source:FlyBase | **-0.56** |
| RPL38 | 3355144 | Ribosomal protein L38 [Source:FlyBase | **-0.51** |
| RPL39 | 37849 | Ribosomal protein L39 [Source:FlyBase | **-0.57** |
| RPS29 | 41200 | Ribosomal protein S29 [Source:FlyBase | **-0.46** |
| SCARNA:MEU5-C46 | 3772530 | small Cajal body-specific RNA : MeU5-C46 [Source:FlyBase | **-0.55** |
| SIRUP | 34089 | Starvation-upregulated protein [Source:FlyBase | **-0.62** |
| SKL | 40016 | sickle [Source:FlyBase | **-0.52** |
| SNORNA:PSI18S-1389A | 5740842 | snoRNA:Psi18S-1389a [Source:FlyBase | **-0.63** |
| SNORNA:PSI28S-1135F | 5740107 | snoRNA:Psi28S-1135f [Source:FlyBase | **-0.58** |
| SNORNA:PSI28S-1175A | 5740342 | snoRNA:Psi28S-1175a [Source:FlyBase | **-0.62** |
| SNORNA:PSI28S-1180 | 3771901 | snoRNA:Psi28S-1180 [Source:FlyBase | **-0.58** |
| SNORNA:PSI28S-2626 | 5740708 | snoRNA:Psi28S-2626 [Source:FlyBase | **-0.62** |
| SNORNA:PSI28S-3342 | 3771886 | snoRNA:Psi28S-3342 [Source:FlyBase | **-0.69** |
| SNRNA:U2:34ABB | 3771927 | small nuclear RNA U2 at 34ABb [Source:FlyBase | **-0.70** |
| SNRNA:U4ATAC:82E | 12798231 | small nuclear RNA U4atac at 82E [Source:FlyBase | **-0.62** |
| SNRNA:U6:96AA | 3772327 | small nuclear RNA U6 at 96Aa [Source:FlyBase | **-0.56** |
| SRA | 47384 | sarah [Source:FlyBase | **-0.31** |
| SU(STE):CR42426 | 7354460 | Su(Ste):CR42426 [Source:FlyBase | **0.68** |
| SU(STE):CR42430 | 7354464 | Su(Ste):CR42430 [Source:FlyBase | **0.63** |
| SULF1 | 53437 | Sulfated [Source:FlyBase | **0.40** |
| TBP | 37476 | TATA binding protein [Source:FlyBase | **-0.45** |
| UGT86DI | 53502 | Ugt86Di [Source:FlyBase | **0.61** |
| ZIP | 38001 | zipper [Source:FlyBase | **0.25** |

**Supplementary Table 5. Differentially expressed genes between 0.4 Gy LDDR + 10 Gy *vs*. 0.4 Gy LDDR irradiated larval male brains.**

A list of 83 differentially expressed genes (72 up-regulated and 11 down-regulated) from 0.4 Gy LDDR + 10 Gy *vs*. 0.4 Gy LDDR (D *vs*. C) comparison. considering a 5% false discovery rate. In the last column are represented the expression levels of each transcript calculated as Log2(D/C).

| **FlyBase ID** | **Entrez ID** | **Description** | **Log2(D/C)** |
| --- | --- | --- | --- |
| agt | 40816 | O-6-alkylguanine-DNA alkyltransferase(agt) | **0.60** |
| AOX1 | 41894 | Aldehyde oxidase 1(AOX1) | **1.22** |
| Bace | 34182 | beta-site APP-cleaving enzyme(Bace) | **-0.35** |
| baz | 32703 | bazooka(baz) | **0.43** |
| CG10005 | 41451 | CG10005 gene product from transcript CG10005-RA(CG10005) | **0.75** |
| CG10445 | 40938 | CG10445 gene product from transcript CG10445-RD(CG10445) | **0.62** |
| CG10570 | 50466 | CG10570 gene product from transcript CG10570-RC(CG10570) | **-0.64** |
| CG10638 | 39424 | CG10638 gene product from transcript CG10638-RD(CG10638) | **0.74** |
| CG11897 | 43450 | CG11897 gene product from transcript CG11897-RB(CG11897) | **1.04** |
| CG12264 | 34613 | CG12264 gene product from transcript CG12264-RA(CG12264) | **0.75** |
| CG15784 | 31461 | CG15784 gene product from transcript CG15784-RB(CG15784) | **1.04** |
| CG1582 | 32021 | CG1582 gene product from transcript CG1582-RB(CG1582) | **0.51** |
| CG17104 | 34465 | CG17104 gene product from transcript CG17104-RB(CG17104) | **0.71** |
| CG1809 | 35981 | CG1809 gene product from transcript CG1809-RA(CG1809) | **-0.01** |
| CG18213 | 42055 | CG18213 gene product from transcript CG18213-RD(CG18213) | **0.65** |
| CG18547 | 41452 | CG18547 gene product from transcript CG18547-RA(CG18547) | **1.36** |
| CG2064 | 35708 | CG2064 gene product from transcript CG2064-RA(CG2064) | **0.58** |
| CG3008 | 33693 | CG3008 gene product from transcript CG3008-RA(CG3008) | **0.88** |
| CG30269 | 37600 | CG30269 gene product from transcript CG30269-RB(CG30269) | **-0.02** |
| CG31869 | 34490 | CG31869 gene product from transcript CG31869-RC(CG31869) | **0.67** |
| CG33158 | 39834 | CG33158 gene product from transcript CG33158-RB(CG33158) | **0.50** |
| CG33502 | 2768875 | CG33502 gene product from transcript CG33502-RA(CG33502) | **0.25** |
| CG3448 | 39094 | CG3448 gene product from transcript CG3448-RB(CG3448) | **0.78** |
| CG42304 | 38332 | CG42304 gene product from transcript CG42304-RA(CG42304) | **0.10** |
| CG43427 | 40583 | CG43427 gene product from transcript CG43427-RP(CG43427) | **0.31** |
| CG43693 | 39231 | CG43693 gene product from transcript CG43693-RD(CG43693) | **-0.40** |
| CG5205 | 41891 | CG5205 gene product from transcript CG5205-RA(CG5205) | **0.57** |
| CG5955 | 40268 | CG5955 gene product from transcript CG5955-RA(CG5955) | **0.76** |
| CG7627 | 34148 | CG7627 gene product from transcript CG7627-RB(CG7627) | **0.64** |
| CG8064 | 42196 | CG8064 gene product from transcript CG8064-RA(CG8064) | **0.29** |
| CHKov2 | 43068 | CG10675 gene product from transcript CG10675-RA(CHKov2) | **0.62** |
| Corp | 31764 | Companion of reaper(Corp) | **0.65** |
| Cpr78E | 40408 | Cuticular protein 78E(Cpr78E) | **-0.76** |
| CR43144 | 19834780 | ncRNA(CR43144) | **1.05** |
| CR44751 | 19834965 | ncRNA(CR44751) | **0.46** |
| Cul2 | 35420 | Cullin 2(Cul2) | **0.54** |
| cv-2 | 45280 | crossveinless 2(cv-2) | **0.59** |
| Cyp6a20 | 36664 | CG10245 gene product from transcript CG10245-RB(Cyp6a20) | **0.61** |
| dally | 39013 | division abnormally delayed(dally) | **0.35** |
| Dif | 35045 | Dorsal-related immunity factor(Dif) | **0.52** |
| dyl | 38531 | dusky-like(dyl) | **0.75** |
| E(spl)m5-HLH | 43158 | Enhancer of split m5. helix-loop-helix(E(spl)m5-HLH) | **-0.57** |
| Ect4 | 38895 | Ectoderm-expressed 4(Ect4) | **0.32** |
| egr | 36054 | eiger(egr) | **0.43** |
| Gadd45 | 35646 | CG11086 gene product from transcript CG11086-RA(Gadd45) | **0.77** |
| Gcn5 | 39431 | Gcn5 ortholog(Gcn5) | **0.41** |
| GstD3 | 48336 | Glutathione S transferase D3(GstD3) | **0.47** |
| GstE6 | 37111 | Glutathione S transferase E6(GstE6) | **0.92** |
| hid | 40009 | head involution defective(hid) | **0.88** |
| Hmgcr | 42803 | HMG Coenzyme A reductase(Hmgcr) | **0.47** |
| Ilp8 | 39909 | Insulin-like peptide 8(Ilp8) | **1.23** |
| Invadolysin | 49580 | CG3953 gene product from transcript CG3953-RA(Invadolysin) | **0.28** |
| Irbp | 117419 | Inverted repeat-binding protein(Irbp) | **0.78** |
| Ku80 | 34930 | CG18801 gene product from transcript CG18801-RA(Ku80) | **0.78** |
| l(3)72Ab | 39737 | lethal (3) 72Ab(l(3)72Ab) | **0.35** |
| lig3 | 41518 | DNA ligase III(lig3) | **0.46** |
| mahj | 37462 | mahjong(mahj) | **0.29** |
| Mocs1 | 39238 | Molybdenum cofactor synthesis 1 ortholog(Mocs1) | **0.60** |
| mre11 | 34565 | meiotic recombination 11(mre11) | **0.64** |
| MRP | 34686 | Multidrug-Resistance like Protein 1(MRP) | **0.76** |
| mus205 | 47186 | mutagen-sensitive 205(mus205) | **0.67** |
| Nplp4 | 50190 | Neuropeptide-like precursor 4(Nplp4) | **0.64** |
| pic | 41611 | piccolo(pic) | **0.35** |
| POSH | 36990 | Plenty of SH3s(POSH) | **0.33** |
| pre-mod(mdg4)-K | 19835678 | CG44879 gene product from transcript CG44879-RB(pre-mod(mdg4)-K) | **-0.08** |
| rad50 | 37564 | CG6339 gene product from transcript CG6339-RE(rad50) | **0.55** |
| RnrL | 34392 | Ribonucleoside diphosphate reductase large subunit(RnrL) | **0.39** |
| RpA-70 | 40972 | Replication Protein A 70(RpA-70) | **0.45** |
| rpr | 40015 | reaper(rpr) | **0.74** |
| scaf | 35505 | scarface(scaf) | **0.61** |
| sda | 44359 | slamdance(sda) | **0.38** |
| Sox14 | 37822 | Sox box protein 14(Sox14) | **-0.46** |
| spn-E | 41919 | spindle E(spn-E) | **1.04** |
| Su(Ste):CR42425 | 7354459 | ncRNA(Su(Ste):CR42425) | **-0.58** |
| Sulf1 | 53437 | Sulfated(Sulf1) | **0.56** |
| thetaTry | 36218 | thetaTrypsin(thetaTry) | **-0.27** |
| Tom | 39619 | Twin of m4(Tom) | **0.45** |
| Traf4 | 33638 | TNF-receptor-associated factor 4(Traf4) | **0.45** |
| Ugt86Da | 53510 | CG18578 gene product from transcript CG18578-RA(Ugt86Da) | **0.83** |
| Ugt86Di | 53502 | CG6658 gene product from transcript CG6658-RB(Ugt86Di) | **1.08** |
| upd2 | 32805 | unpaired 2(upd2) | **0.58** |
| wnd | 40143 | wallenda(wnd) | **0.29** |
| zip | 38001 | zipper(zip) | **0.31** |

**Supplementary Table 6. Differentially expressed genes between 10 Gy-irradiated *vs*. unirradiated larval male brains.**

A list of 82 differentially expressed genes (42 up-regulated and 40 down-regulated) from 10 Gy *vs*. unirradiated (B *vs*. A) comparison. considering a 5% false discovery rate. In the last column are represented the expression levels of each transcript calculated as Log2(B/A).

| **FlyBase ID** | **Entrez ID** | **Description** | **log2(B/A)** |
| --- | --- | --- | --- |
| Aldh | 34256 | Aldehyde dehydrogenase(Aldh) | **0.49** |
| AOX1 | 41894 | Aldehyde oxidase 1(AOX1) | **0.85** |
| AstC | 34537 | Allatostatin C(AstC) | **0.42** |
| bbx | 3772670 | bobby sox(bbx) | **0.37** |
| Brca2 | 37916 | Breast cancer 2. early onset homolog(Brca2) | **-0.40** |
| CG11029 | 33804 | CG11029 gene product from transcript CG11029-RA(CG11029) | **-0.42** |
| CG11147 | 33803 | CG11147 gene product from transcript CG11147-RA(CG11147) | **-0.36** |
| CG11897 | 43450 | CG11897 gene product from transcript CG11897-RB(CG11897) | **0.55** |
| CG12539 | 32412 | CG12539 gene product from transcript CG12539-RA(CG12539) | **-0.56** |
| CG1287 | 40898 | CG1287 gene product from transcript CG1287-RA(CG1287) | **-0.66** |
| CG13067 | 39785 | CG13067 gene product from transcript CG13067-RA(CG13067) | **0.97** |
| CG13631 | 50073 | CG13631 gene product from transcript CG13631-RB(CG13631) | **0.53** |
| CG15784 | 31461 | CG15784 gene product from transcript CG15784-RB(CG15784) | **0.69** |
| CG17680 | 37071 | CG17680 gene product from transcript CG17680-RA(CG17680) | **0.37** |
| CG18547 | 41452 | CG18547 gene product from transcript CG18547-RA(CG18547) | **1.22** |
| CG1882 | 35733 | CG1882 gene product from transcript CG1882-RG(CG1882) | **0.50** |
| CG30046 | 36334 | CG30046 gene product from transcript CG30046-RC(CG30046) | **-0.54** |
| CG3008 | 33693 | CG3008 gene product from transcript CG3008-RA(CG3008) | **0.47** |
| CG31921 | 319027 | CG31921 gene product from transcript CG31921-RA(CG31921) | **-0.56** |
| CG32365 | 38876 | CG32365 gene product from transcript CG32365-RB(CG32365) | **0.47** |
| CG32500 | 2768879 | CG32500 gene product from transcript CG32500-RA(CG32500) | **0.56** |
| CG34232 | 5740195 | CG34232 gene product from transcript CG34232-RC(CG34232) | **0.54** |
| CG3448 | 39094 | CG3448 gene product from transcript CG3448-RB(CG3448) | **0.59** |
| CG4096 | 31490 | CG4096 gene product from transcript CG4096-RB(CG4096) | **0.41** |
| CG43192 | 12798442 | CG43192 gene product from transcript CG43192-RA(CG43192) | **-0.64** |
| CG5080 | 33299 | CG5080 gene product from transcript CG5080-RB(CG5080) | **0.52** |
| CG6191 | 36513 | CG6191 gene product from transcript CG6191-RB(CG6191) | **0.39** |
| CG8771 | 36397 | CG8771 gene product from transcript CG8771-RB(CG8771) | **-0.24** |
| cher | 42066 | cheerio(cher) | **0.39** |
| Cht6 | 31935 | CG43374 gene product from transcript CG43374-RK(Cht6) | **0.51** |
| ci | 43767 | cubitus interruptus(ci) | **-0.41** |
| conu | 3355133 | conundrum(conu) | **-0.38** |
| Corp | 31764 | Companion of reaper(Corp) | **0.60** |
| CR33987 | 3885608 | ncRNA(CR33987) | **0.63** |
| CR44017 | 14462439 | ncRNA(CR44017) | **-0.63** |
| CR44161 | 14462548 | ncRNA(CR44161) | **-0.70** |
| CR44993 | 19835215 | ncRNA(CR44993) | **-0.55** |
| CR45137 | 19835272 | ncRNA(CR45137) | **-0.53** |
| CR45820 | 26067148 | ncRNA(CR45820) | **-0.54** |
| CR45993 | 26067314 | ncRNA(CR45993) | **-0.75** |
| CR46003 | 26067324 | ncRNA(CR46003) | **-0.33** |
| cv-2 | 45280 | crossveinless 2(cv-2) | **0.40** |
| Cyp310a1 | 35115 | CG10391 gene product from transcript CG10391-RA(Cyp310a1) | **-0.58** |
| Cyp4ac2 | 33755 | CG17970 gene product from transcript CG17970-RB(Cyp4ac2) | **-0.55** |
| fat-spondin | 36919 | CG6953 gene product from transcript CG6953-RA(fat-spondin) | **0.45** |
| Gadd45 | 35646 | CG11086 gene product from transcript CG11086-RA(Gadd45) | **0.75** |
| GstT3 | 33047 | Glutathione S transferase T3(GstT3) | **0.53** |
| GstZ2 | 41133 | Glutathione S transferase Z2(GstZ2) | **-0.32** |
| hid | 40009 | head involution defective(hid) | **0.77** |
| Ilp8 | 39909 | Insulin-like peptide 8(Ilp8) | **0.44** |
| Irbp | 117419 | Inverted repeat-binding protein(Irbp) | **0.52** |
| JhI-26 | 36819 | Juvenile hormone-inducible protein 26(JhI-26) | **0.76** |
| kra | 40680 | krasavietz(kra) | **-0.34** |
| l(2)k09022 | 33960 | lethal (2) k09022(l(2)k09022) | **-0.33** |
| magu | 36048 | CG2264 gene product from transcript CG2264-RA(magu) | **-0.48** |
| MRP | 34686 | Multidrug-Resistance like Protein 1(MRP) | **0.34** |
| myo | 43811 | myoglianin(myo) | **-0.35** |
| Npc1a | 34358 | Niemann-Pick type C-1a(Npc1a) | **-0.24** |
| Nrx-IV | 39387 | Neurexin IV(Nrx-IV) | **-0.29** |
| Osi14 | 40770 | Osiris 14(Osi14) | **0.57** |
| pan | 43769 | pangolin(pan) | **-0.41** |
| path | 39106 | pathetic(path) | **-0.48** |
| Pten | 43991 | Phosphatase and tensin homolog(Pten) | **-0.43** |
| Pvf2 | 33994 | PDGF- and VEGF-related factor 2(Pvf2) | **0.50** |
| Qtzl | 318990 | Quetzalcoatl(Qtzl) | **0.62** |
| roX1 | 3772376 | RNA on the X 1(roX1) | **-0.50** |
| rpr | 40015 | reaper(rpr) | **0.67** |
| RpS9 | 39108 | Ribosomal protein S9(RpS9) | **-0.30** |
| Rpt3R | 41190 | Regulatory particle triple-A ATPase 3-related(Rpt3R) | **0.56** |
| scf | 38145 | supercoiling factor(scf) | **0.37** |
| Sema-2b | 246538 | Semaphorin-2b(Sema-2b) | **-0.36** |
| snRNA:U2:34ABb | 3771927 | small nuclear RNA U2 at 34ABb(snRNA:U2:34ABb) | **0.62** |
| Sox102F | 43844 | CG11153 gene product from transcript CG11153-RB(Sox102F) | **-0.36** |
| spok | 5740359 | spookier(spok) | **-0.42** |
| Stam | 34505 | Signal transducing adaptor molecule(Stam) | **0.27** |
| Su(Ste):CR42410 | 7354443 | ncRNA(Su(Ste):CR42410) | **-0.70** |
| Su(Ste):CR42415 | 7354448 | ncRNA(Su(Ste):CR42415) | **-0.66** |
| Su(Ste):CR42426 | 7354460 | ncRNA(Su(Ste):CR42426) | **-0.75** |
| Su(Ste):CR42430 | 7354464 | ncRNA(Su(Ste):CR42430) | **-0.72** |
| Syp | 42460 | Syncrip(Syp) | **-0.30** |
| Tom | 39619 | Twin of m4(Tom) | **0.42** |
| Xrp1 | 42267 | CG17836 gene product from transcript CG17836-RB(Xrp1) | **0.50** |

**Supplementary Table 7. Differentially expressed genes between 10 Gy *vs*. 0.4 Gy LDDR irradiated larval male brains.**

A list of 52 differentially expressed genes (31 up-regulated and 21 down-regulated) from 10 Gy *vs*. 0.4 Gy LDDR (B *vs*. C) comparison. considering a 5% false discovery rate. In the last column are represented the expression levels of each transcript calculated as Log2(B/C).

| **FlyBase ID** | **ENTREZ ID** | **Description** | **Log2(B/C)** |
| --- | --- | --- | --- |
| ana | 35913 | anachronism | **0.70** |
| AOX1 | 41894 | Aldehyde oxidase 1 | **0.92** |
| CG13067 | 39785 | uncharacterized protein | **1.36** |
| CG13403 | 32366 | uncharacterized protein | **-3.85** |
| CG13460 | 39638 | uncharacterized protein | **-1.17** |
| CG1468 | 31920 | uncharacterized protein | **1.26** |
| CG15034 | 31670 | uncharacterized protein | **-2.51** |
| CG15784 | 31461 | uncharacterized protein | **0.91** |
| CG18547 | 41452 | uncharacterized protein | **1.52** |
| CG31789 | 318943 | uncharacterized protein | **-5.81** |
| CG32198 | 317909 | uncharacterized protein | **-2.54** |
| CG32447 | 40412 | uncharacterized protein | **-0.50** |
| CG3819 | 40069 | uncharacterized protein | **-4.23** |
| CG42326 | 35838 | uncharacterized protein | **0.84** |
| CG42516 | 8674068 | uncharacterized protein | **-5.68** |
| CG5080 | 33299 | uncharacterized protein | **0.58** |
| CG5955 | 40268 | uncharacterized protein | **0.66** |
| CG6839 | 40067 | uncharacterized protein | **-4.24** |
| cher | 42066 | cheerio | **0.39** |
| Cht10 | 3355116 | Chitinase 10 | **-0.74** |
| Cyp28d1 | 33749 | Cyp28d1 | **0.62** |
| Fbp1 | 39566 | Fat body protein 1 | **2.85** |
| Gadd45 | 35646 | Growth arrest and DNA damage-inducible 45 | **0.90** |
| hid | 40009 | head involution defective | **0.95** |
| IBIN | 19835457 | Induced by Infection | **2.40** |
| Ilp8 | 39909 | Insulin-like peptide 8 | **1.61** |
| Irbp | 117419 | Inverted repeat-binding protein | **0.65** |
| lncRNA:CR43144 | 19834780 | long non-coding RNA:CR43144 | **1.37** |
| LysB | 38125 | Lysozyme B | **-6.82** |
| LysD | 38127 | Lysozyme D | **-4.96** |
| magu | 36048 | magu | **-0.55** |
| mam | 36555 | mastermind | **-1.42** |
| MRP | 34686 | Multidrug-Resistance like Protein 1 | **0.37** |
| Muc68E | 2768990 | Mucin 68E | **-2.76** |
| Obp99b | 43497 | Odorant-binding protein 99b | **1.85** |
| obst-A | 33022 | obstructor-A | **-0.71** |
| path | 39106 | pathetic | **-0.69** |
| Phae1 | 34637 | Phaedra 1 | **-3.50** |
| Picot | 36865 | picot | **0.55** |
| Pig1 | 31303 | Pre-intermoult gene 1 | **-1.31** |
| rdog | 43450 | red dog mine | **0.78** |
| rpr | 40015 | reaper | **0.84** |
| spn-E | 41919 | spindle E | **0.74** |
| Su(Ste):CR42407 | 7354440 | ncRNA | **-1.98** |
| Su(Ste):CR42410 | 7354443 | ncRNA | **-1.58** |
| Su(Ste):CR42415 | 7354448 | ncRNA | **-1.60** |
| Su(Ste):CR42424 | 7354458 | ncRNA | **-2.21** |
| Su(Ste):CR42425 | 7354459 | ncRNA | **-1.37** |
| Su(Ste):CR42426 | 7354460 | ncRNA | **-1.93** |
| Su(Ste):CR42430 | 7354464 | ncRNA | **-1.66** |
| vir-1 | 34652 | virus-induced RNA 1 | **0.57** |
| Wnt2 | 35975 | Wnt oncogene analog 2 | **-0.65** |

**Supplementary Table 8 RNA processing biological process**.

A list of 18 differentially expressed genes between 0.4 Gy LDDR + 10 Gy and 10 Gy (D *vs*. B comparison) that are enriched in “RNA processing” biological process obtained from DAVID functional analysis (Adjusted p-value < 0.05). For each sample is represented the normalized expression value of each gene.

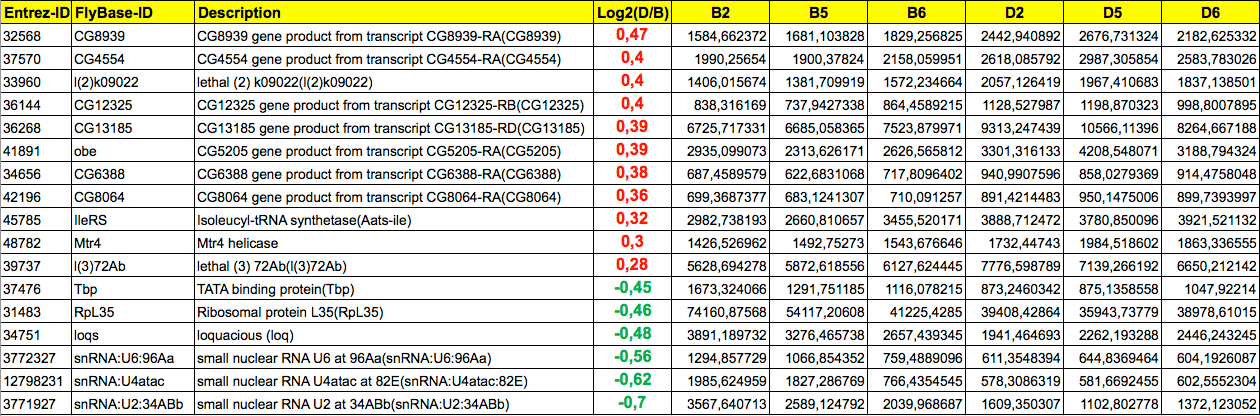
